# Determining the stages of cellular differentiation using Deep Ultraviolet Resonance Raman Spectroscopy

**DOI:** 10.1101/2020.11.08.373001

**Authors:** Nicole M. Ralbovsky, Paromita Dey, Bijan K. Dey, Igor K. Lednev

## Abstract

Cellular differentiation is a fundamental process in which one cell type changes into one or more specialized cell types. Cellular differentiation starts at the beginning of embryonic development when a simple zygote begins to transform into a complex multicellular organism composed of various cell and tissue types. This process continues into adulthood when adult stem cells differentiate into more specialized cells for normal growth, regeneration, repair, and cellular turnover. Any abnormalities associated with this fundamental process of cellular differentiation is linked to life threatening conditions including degenerative diseases and cancers. Detection of undifferentiated and different stages of differentiated cells can be used for disease diagnosis but is often challenging due to the laborious procedures, expensive tools, and specialized technical skills which are required. Here, a novel approach, called deep ultraviolet resonance Raman spectroscopy, is used to study various stages of cellular differentiation using a well-known myoblast cell line as a model system. These cells proliferate in the growth medium and spontaneously differentiate in differentiation medium into myocytes and later into myotubes and myofibers. The cellular and molecular characteristics of these cells mimic very well actual muscle tissue *in vivo*. We have found that undifferentiated myoblast cells and myoblast cells differentiated at three different stages are able to be easily separated using deep ultraviolet resonance Raman spectroscopy in combination with chemometric techniques. Our study has a great potential to study cellular differentiation during normal development as well as to detect abnormal cellular differentiation in human pathological conditions in future studies.

## Introduction

Within multicellular organisms, tissues are organized as a collection of cells which differentiate from totipotent fertilized embryos to carry out specific physiological functions. The balance between cellular proliferation and differentiation is critical for normal physiological function and health. Disruption of this balance is associated with numerous human conditions including degenerative diseases(1) and cancer(2). In this study, a skeletal muscle stem cell (MuSCs)-derived myoblast cell line is used as a model system. Postnatal skeletal muscle development, growth, regeneration, and maintenance of homeostasis depends on MuSCs, also known as satellite cells. MuSCs reside beneath the basal lamina juxtaposed to the muscle fiber and are mitotically quiescent. In response to muscle injury, quiescent MuSCs are activated to reenter the cell cycle, followed by proliferation to form a pool of myoblasts, and eventually exit at the G1 phase in the cycle to then differentiate and fuse into newly formed or existing myofibers. A subset of MuSCs are self-renewed and return to quiescence. This extensive process of making new muscle fibers is known as myogenesis and is quintessential for normal physiological function. If anything goes awry in these processes at the cellular or molecular level, human diseases like Duchenne muscular dystrophy or soft tissue cancer, called rhabdomyosarcoma, can arise. The gene expression program, including transcription factors and signaling molecules that govern myogenesis, has been well characterized. However, the processes involved to determine the stages of cellular differentiation through measurement of these molecular signatures are laborious and expensive and require specialized skills to implement. Thus, a new method for achieving this goal was explored using deep ultraviolet resonance Raman spectroscopy (DUVRS).

The advantages of DUVRS make it a suitable method for exploring various biological specimens and phenomena. DUVRS has been used in the past for investigating malignant biological specimens(3), respiratory diseases(4), and for studying protein structure and transformation(5-7) as well examining protein aggregates and fibrillogenesis.(8-11) Excitation in the deep ultraviolet (UV) range is known to enhance the inelastic scattering of many biological samples.(12) Specifically, the Raman signal of polypeptide side chains including aromatic amino acids are strongly resonantly enhanced. Aromatic amino acids such as tryptophan and tyrosine strongly absorb UV light around 280 nm and 230 nm which allows for the resonance enhancement of their Raman scattering.(12) Strong resonance enhancement of Raman scattering from phenylalanine occurs at deep UV excitation below 200 nm.(10) Resonance Raman spectra of aromatic amino acid residues provide important information about the tertiary structure of proteins. Additionally, deep UV excitation resonantly enhances the Raman scattering of the amide chromophore, a building block of the polypeptide backbone.(13) This enhancement provides information regarding the secondary structure of a protein; as such, DUVRS is extremely useful for investigating proteins within biological samples. Resonance enhancement of nucleic acids has additionally been observed via deep UV excitation due to their absorption of light in the same range.(14, 15) Both proteins and nucleic acids play influential roles in biochemical processes and are therefore anticipated to be useful for distinguishing between biological samples, making deep UV excitation uniquely advantageous when compared to excitation using visible or near-IR light.

Along with providing unique enhancement of signals from crucial biomolecules, DUVRS typically produces a stronger signal-to-noise ratio in the resultant Raman spectrum due to the absence of fluorescence interference.(16) Fluorescence typically occurs at wavelengths longer than 250 nm, thus shifting the Raman excitation wavelength to be shorter than 250 nm will allow for a much better quality spectrum to obtained due to the lack of fluorescence interference.(12) A better signal-to-noise ratio is crucial for examining biological samples such as cells in a liquid suspension. Typical Raman excitation in the visible or near-IR range will not produce the same quality of spectrum due to strong fluoresce accompanied with such a sample.

DUVRS is used here to investigate various stages of myoblast differentiation. Results show that all four stages which were studied were successfully discriminated from each other using chemometric analysis. These results indicate the potential of the method to study abnormal and/or differential cellular differentiation in human pathological conditions including cancer, such as rhabdomyosarcoma, that arise from abnormal myoblast differentiation.(2)

## Materials and Methods

### Myoblast cell culture and differentiation assay

Mouse myoblast cell line (C2C12) was acquired from the American Type Culture Collection (ATCC; Manassas, VA, USA). Cells were maintained at subconfluent densities in growth medium (GM) at 37 °C in a tissue culture incubator with a constant supply of 5% CO2. GM was made up of Dulbecco’s modified Eagle medium (DMEM; Life Technologies, Carlsbad, CA, USA) supplemented with 10% FBS and 1X antibiotic-antimycotic (Life Technologies).(17, 18) For myogenic differentiation assays, the C2C12 myoblast cells were grown to about 75% confluency, washed with 1X phosphate-buffered saline (PBS), and cultured with differentiation medium (DM). DM was made up of DMEM containing 2% heat-inactivated horse serum (HyClone) and 1X antibiotic-antimycotic (Life Technologies).(17, 18) Cells were harvested while growing in GM and after 48 hours (DM2), 96 hours (DM4) and 144 hours (DM6) in DM. The images of undifferentiated (GM) and different stages of differentiated (DM2, DM4 and DM6) samples were taken using EVOS Cell Imaging Systems (Thermo Fisher Scientific, Waltham, MA, USA).

### Total RNA isolation and quantitative reverse transcription polymerase chain reaction (qRT-PCR) assays

Total RNA was extracted using RNEasy mini kit (Qiagen, Hilden, Germany) by following the manufacturer’s instructions. cDNA synthesis was carried out using the iScript cDNA Synthesis Kit (Bio-Rad, Hercules, CA, USA) as instructed. Then, qRT-PCR was carried out using Sybr green PCR master mix (Bio-Rad) in a Bio-Rad thermal cycler using Myogenin and Myosin Heavy Chain (MHC) specific primers. GAPDH primer pairs were used as a housekeeping gene for normalizing the values of Myogenin and MHC.

### DUVRS analysis of myoblasts

A total of 31 cell samples were analyzed from four different stages of myoblast cell differentiation, including undifferentiated cells (GM, n=8) and cells allowed to differentiate in DM for 48 (DM2, n=8), 96 (DM4, n=8), and 144 hours (DM6, n=7). All samples were analyzed using a custom-built deep ultraviolet Raman spectrograph (details of which can be found elsewhere).(19) Briefly, the samples were excited using 198-nm radiation generated at the 5^th^ anti-Stokes shift from the third harmonic of a Ni-YAG laser in a Raman shifter which is filled with low pressure hydrogen. A UV laser beam (at a power of about 0.5 mW at the surface of the sample) was focused within a spinning Suprasil NMR tube which contained approximately 200 µL of sample solution. The solution was kept continuously spinning with a magnetic stir bar to prevent burning of the sample. Scattered radiation was collected in the backscattering geometry, dispersed via a double monochromator, and detected using a liquid-nitrogen cooled CCD camera.

To acquire the DUVRS spectral data, 20 accumulations of 30 s each were collected per sample. Each accumulation was saved as an individual spectrum to obtain multiple spectra per sample to use for statistical analysis. A comparison was made between the individual spectra acquired for each sample; no gradual changes to the spectra with respect to accumulation number were observed, indicating sample photodegradation due to UV radiation did not occur.

### DUVRS data analysis

620 spectra were obtained from all samples and loaded into GRAMS v9.2 software (Thermo Fisher Scientific). The spectral signature of the quartz NMR tube and of the buffer solution was subtracted from each spectrum individually. Spectra were then calibrated from pixels to wavenumbers using the DUVRS spectrum of Teflon as a standard.

### Chemometric analysis

PLS_Toolbox (Eigenvector Research Inc., Wenatchee, WA, USA) operating within MATLAB software (version 2017b, Mathworks, Inc, Natick, MA, USA) was used for chemometric analysis. Initially, preprocessing steps were performed including spectral smoothing, baseline correction, and normalization. Following data processing, various chemometric methods were applied for distinguishing between the four stages of myoblast cell differentiation. The samples were split into two different datasets: a calibration dataset (n=27) and a validation dataset (n=4). The goal of the analysis was to separate all four stages of myoblast differentiation. Here, genetic algorithm (GA) was applied to reduce the complexity of the spectral dataset and to identify which features were the most useful for discrimination. Then, partial least squares discriminant analysis (PLS-DA) was performed using the GA-identified spectral dataset for building the quaternary model for classification purposes. The performance of the model was evaluated using the donors from the validation dataset.

## Results and Discussion

The C2C12 myoblast cell line serves as an excellent model system for studying cellular differentiation. Differentiation of myoblast cells into myocytes, myotubes, or myofiber-like structures can be achieved in cell culture by reducing serum supplements. As shown in Figure 1, C2C12 myoblast cells proliferate in growth medium (GM) and differentiate in differentiation medium (DM). As the differentiation process progressed, myogenic markers, including Myogenin and MHC, were upregulated. We harvested 8 undifferentiated (GM), DM2, and DM4 samples each and 7 DM6 samples.

**Figure 1.**
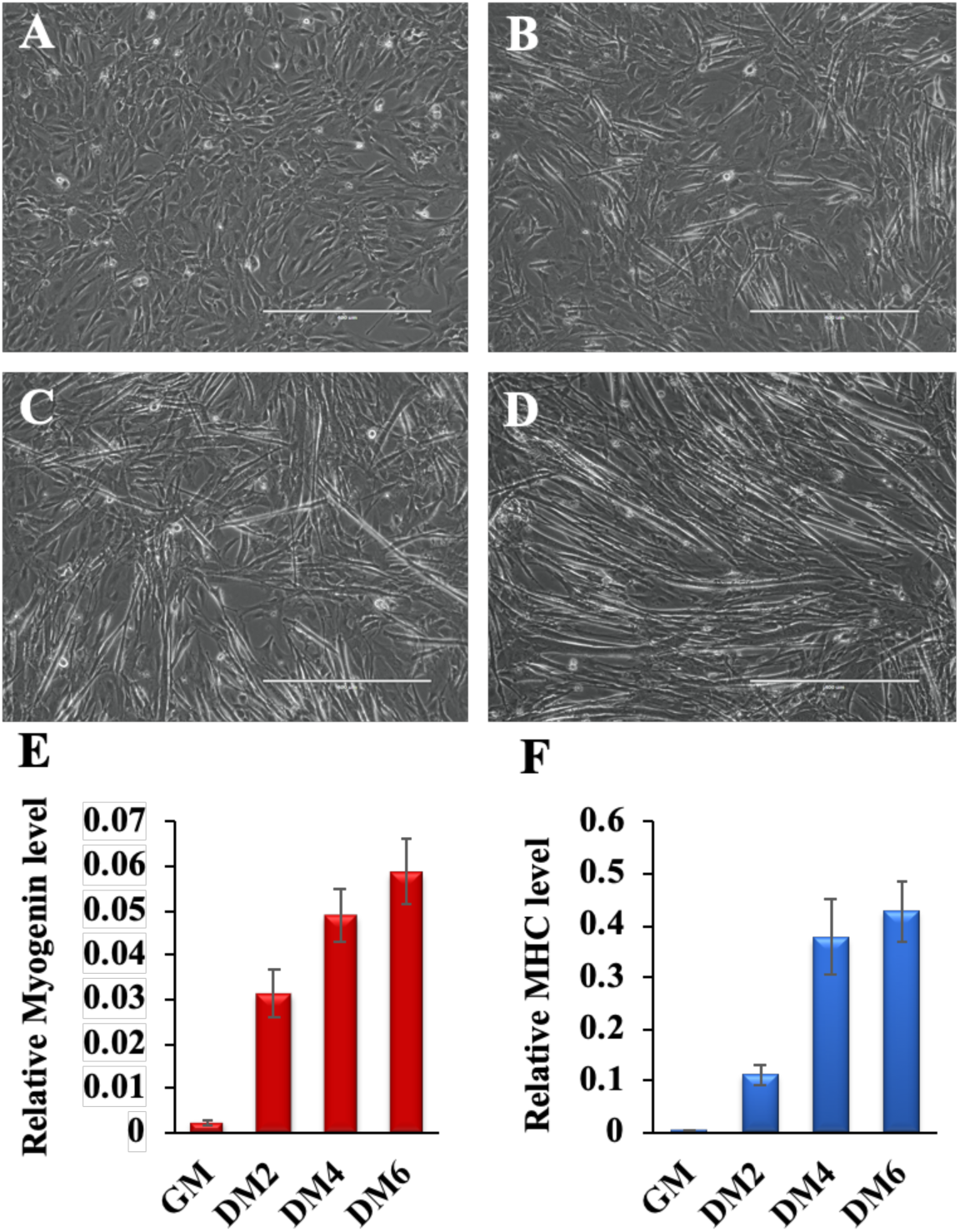
Myoblast cells (C2C12) proliferate in the growth medium and spontaneously differentiate when transferred to differentiation medium. (A) Undifferentiated and (B-D) different stages of differentiated myoblast cells are shown. (B) During early differentiation myoblast cells elongate and differentiate into myocytes and later (C) multiple myocytes fuse together to form myotubes, and subsequently (D) multiple myotubes align together to form myofiber-like structures. (E) An early differentiation marker, Myogenin and (F) a late myogenic marker, myosin heavy chain (MHC) mRNA levels are shown. Myogenin and MHC levels were normalized to GAPDH. As differentiation continues, both Myogenin and MHC expression levels are upregulated. All experiments were done with at least three or more biological replicates. Scale Bar: 400 µM.

A total of 620 DUVRS spectra were collected from the 31 samples. The average spectrum for each of the four classes (GM, DM2, DM4, and DM6) is seen in Figure 2. Each spectrum is the average of all spectra collected from all samples in each class.

**Figure 2.**
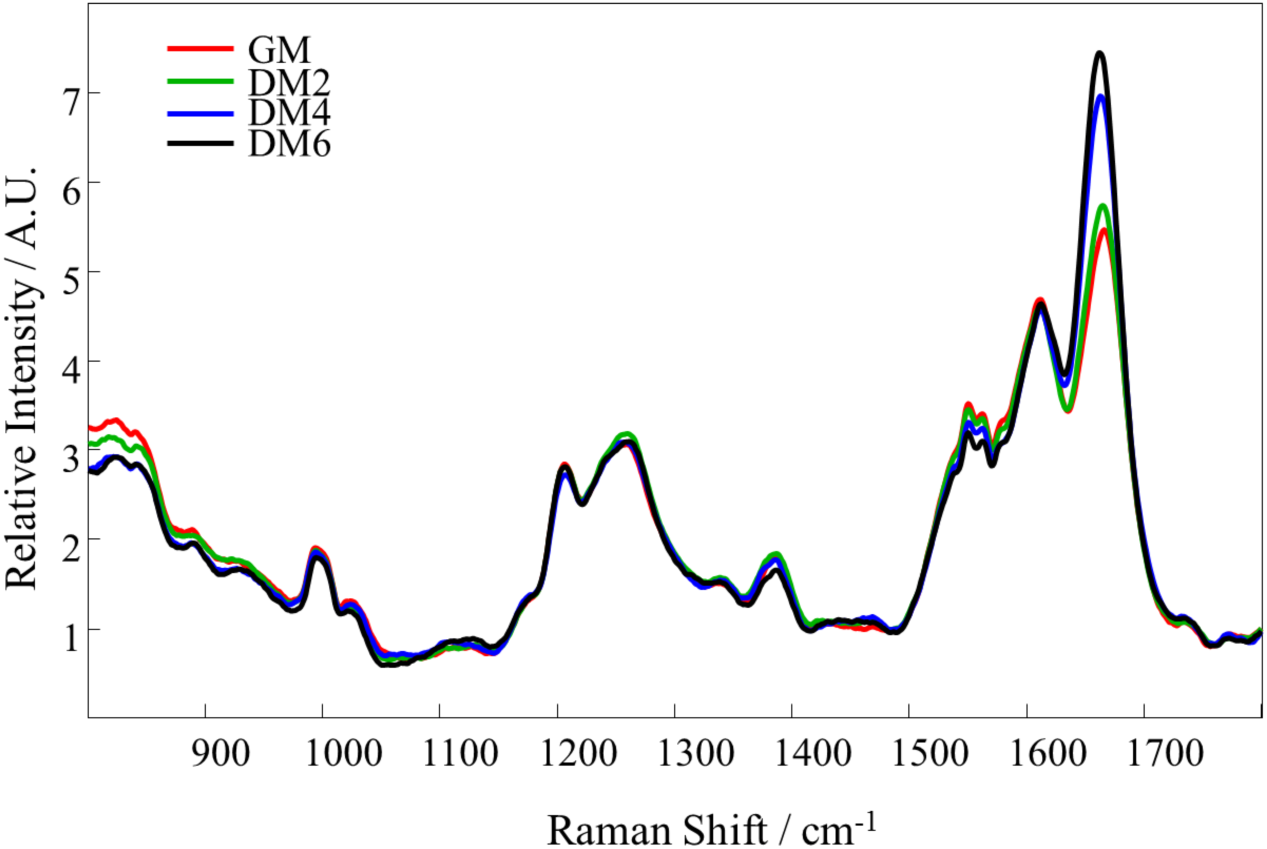
Average Raman spectra obtained for all samples of each of the four stages of myoblast differentiation including undifferentiated cells (GM, red) and cells allowed to differentiate for 48 (DM2, green), 96 (DM4, blue), or 144 hours (DM6, black).

The average spectra appear very similar to each other – this is not surprising due to the anticipated high level of overlap in molecular composition between the differentiation stages. The majority of the peaks which contribute to the spectra correspond to proteins and nucleic acids. A summary of the main peaks and their tentative assignments is given in Table 1.

**Table 1.**
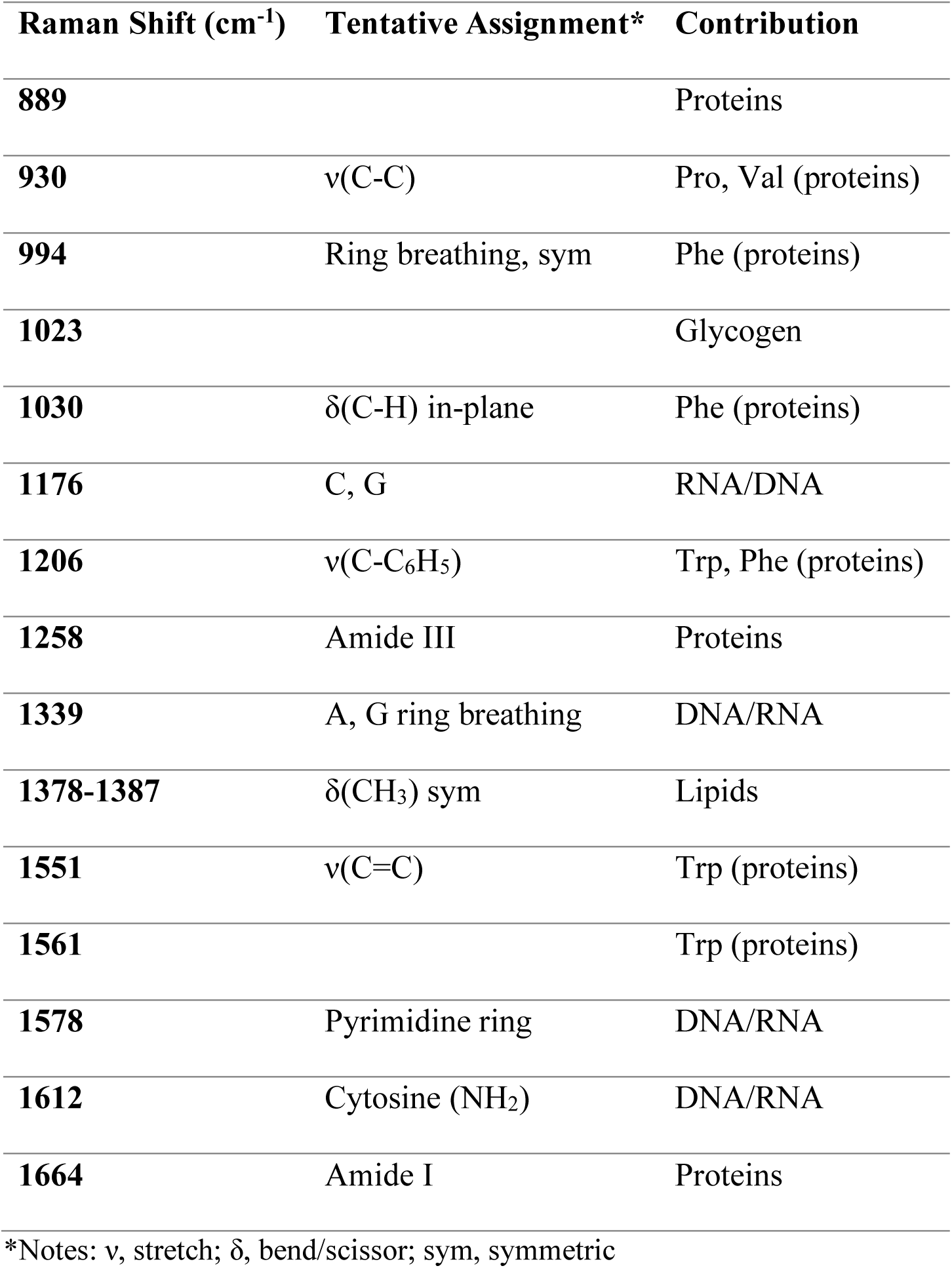
Tentative assignments of the main peaks in the average Raman spectra of myoblasts at progressive stages of differentiation(20)

### Analysis of all four stages of myoblast differentiation

The small changes which are observed between the average spectra of the different myoblasts are not found to be significant. The largest observable difference is seen at 1633 cm^-1^, however when the differences in average intensities at each stage are compared with ±1 standard deviations, the average intensity is not found to be significantly different between the groups (Figure 3). The same is found for other small changes in intensity, including the Raman peaks at 1563 cm^-1^ and 1561 cm^-1^ (supplementary information Figures S.1 and S.2).

**Figure 3.**
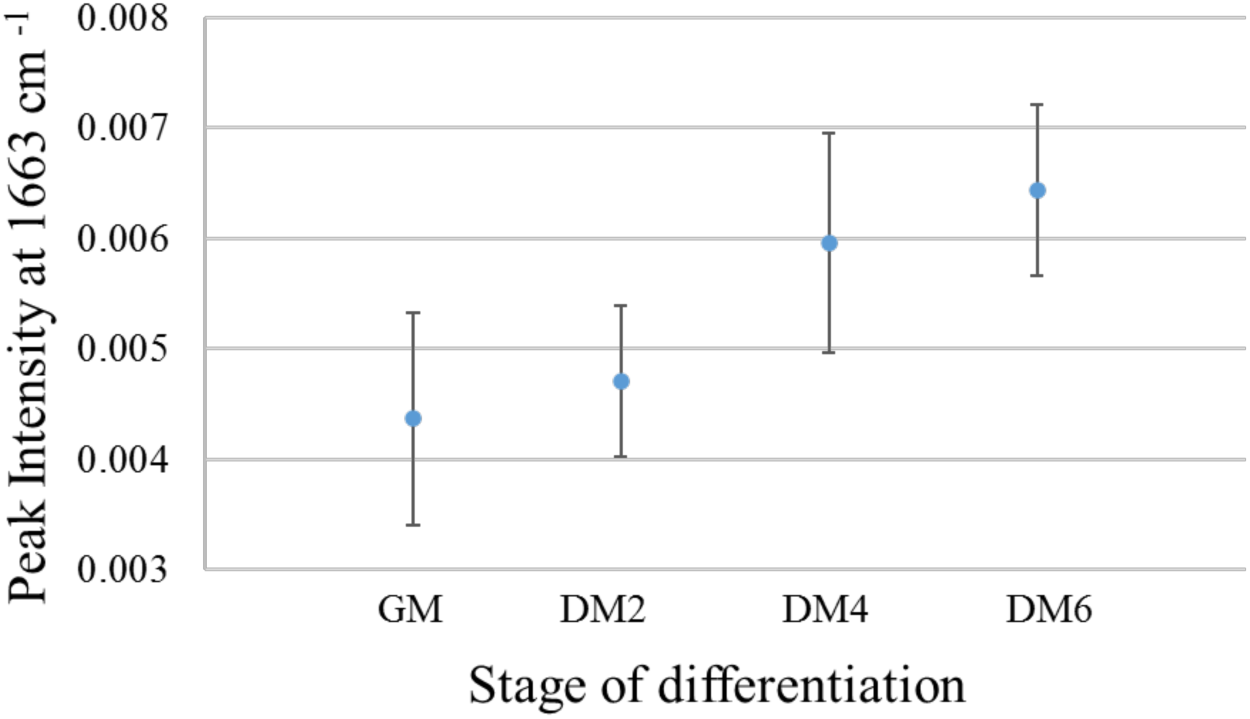
The mean ±1 standard deviation of the 1633 cm^-1^ Raman peak intensity for each of the four classes, demonstrating insignificant differences as observed by the overlap in standard deviations between groups.

Statistical analysis was thus required to discriminate between the four stages of myoblast differentiation. Genetic algorithm was first employed to identify the subset of spectral features which were the most useful for distinguishing between the classes of data and which will support the prediction algorithm’s capabilities. Results of GA (supplementary information, Figure S.3) indicated that proteins and DNA/RNA are the most influential biochemical components which allow for discriminating between the four stages. Myoblast differentiation is a dynamic and robust process; this process shows vigorous changes in a large number of gene expressions, which have been documented in numerous studies during myoblast differentiation.(21-25) Although gene amplification usually occurs in cancer cells, a number of gene amplifications have been reported to occur during myoblast differentiation.(26) The global gene expression has found the changes in a large number of genes during myoblast differentiation shows.(21, 24, 25) Proteomic studies have further confirmed that a large number of proteins are up-regulated in the differentiated myoblast cells.(22, 23) Our own findings from RNA-seq data analysis show that more than four thousand genes are upregulated in the differentiated myotubes, and a subset of pro-myogenic genes such as Casq1, Myh3, Myh4, Actn2 are upregulated more than 2000-fold in these samples (data not shown). Therefore, it is not surprising to see that our findings detect and can discriminate the various stages of differentiated samples based on contributions from DNA/RNA and proteins.

Further, PLS-DA was performed using the GA-identified spectral dataset to build a discriminatory model. The 27 samples of the calibration dataset were used to build the algorithm, and the four samples of the validation dataset were set aside for independent external validation of the method. The model was built using three latent variables, which captured the maximum covariance between the groups. Through internal cross-validation, the model obtained an average accuracy of 75% for correctly predicting the class of a spectrum.

The spectral data of the four independent donors of the validation dataset were then loaded into the model. The spectral predictions generated for the four donors of the calibration dataset indicated 83% accuracy for external validation (Table 2, left panel). Using the spectral-level predictions, overall sample-level predictions were made; here, 100% accuracy was achieved for correctly predicting the stage of differentiation at the sample-level for each of the four samples used for external validation (Table 2, right panel).

**Table 2.** Classification predictions for all individual spectra (left) and overall sample-level classification predictions (right) of the four independent samples used for external validation

| Individual Spectral Predictions |  |  |  |  | External Validation Results |  |  |
| --- | --- | --- | --- | --- | --- | --- | --- |
| *Sample* | *Predicted Class* |  |  |  | *Sample* | *Predicted Class* | *True Class* |
|  | **GM** | **DM2** | **DM4** | **DM6** |  |  |  |
| **#1** | 16 | 4 | 0 | 0 | **#1** | GM | GM |
| **#2** | 4 | 16 | 0 | 0 | **#2** | DM2 | DM2 |
| **#3** | 0 | 4 | 14 | 2 | **#3** | DM4 | DM4 |
| **#4** | 0 | 0 | 7 | 13 | **#4** | DM6 | DM6 |

An additional classification system was generated in a similar manner to discriminate between early (GM and DM2) and late (DM4 and DM6) stages of myoblast cell differentiation in a binary model; this model showed similar levels of success for class separation (supplementary information Figure S.4)

The minimal changes in biochemical composition which occur are shown here to be sufficient for discriminating between four different stages of myoblast differentiation. DNA/RNA and proteins are indicated as the most significant classes of biomolecules for successful separation of the four stages. Importantly, the results obtained during external validation support the capability of the method for classifying spectral data from samples which were not used to build it. This indicates the potential of the method to be expanded upon in the future for clinical applications, such as for the analysis of abnormal cellular differentiation in human pathological conditions including cancer and Duchenne muscular dystrophy. Conducting differentiation at the single cell level using spontaneous Raman spectroscopy could also be possible, as the method was recently reported for Celiac disease diagnostics based on analysis of a single red blood cell.(27)

Deep ultraviolet resonance Raman spectroscopy (DUVRS) is capable of capturing vital information regarding a biological samples’ composition, including information regarding protein structure and nucleic acid composition. DUVRS is used in this study to successfully distinguish between myoblasts which were either undifferentiated or allowed to differentiate for varying numbers of hours. Specifically, a model was built using GA and PLS-DA for distinguishing between four stages of myoblast differentiation. This model achieved 100% successful classification at the level of individual sample during external validation. Analysis of the DUVRS spectra indicate that biochemical changes which occur during cell differentiation stem mostly from proteins and nucleic acids. DUVRS is fully capable of discriminating between the various stages of myoblast differentiation, opening the door for future exploration into cellular differentiation during normal development as well as into detecting abnormal cellular differentiation in human pathological conditions.

## Acknowledgments

This work was supported by the SUNY startup, the American Heart Association (AHA 17SDG33670339) and the National Institute of Arthritis and Musculoskeletal and Skin Diseases, NIAMS (R15AR074728) grants to B.K.D. N.M.R was supported by NIH training Grant T32 GM13206.

## Supplementary Information

**Figure S.1.**
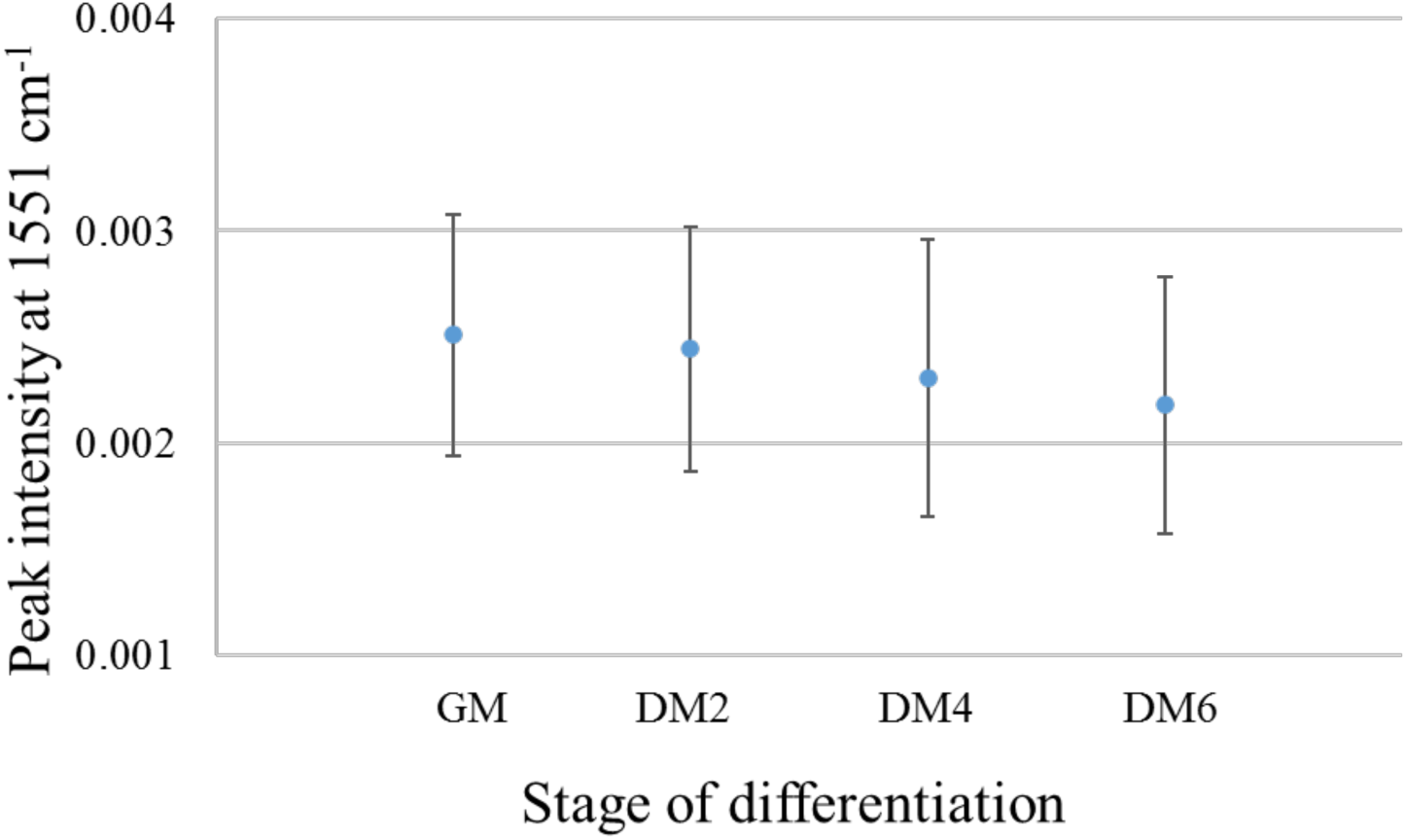
The mean ±1 standard deviation of the 1551 cm^-1^ peak intensity for each of the four classes, demonstrating insignificant differences as observed by the overlap in standard deviations between groups.

**Figure S.2.**
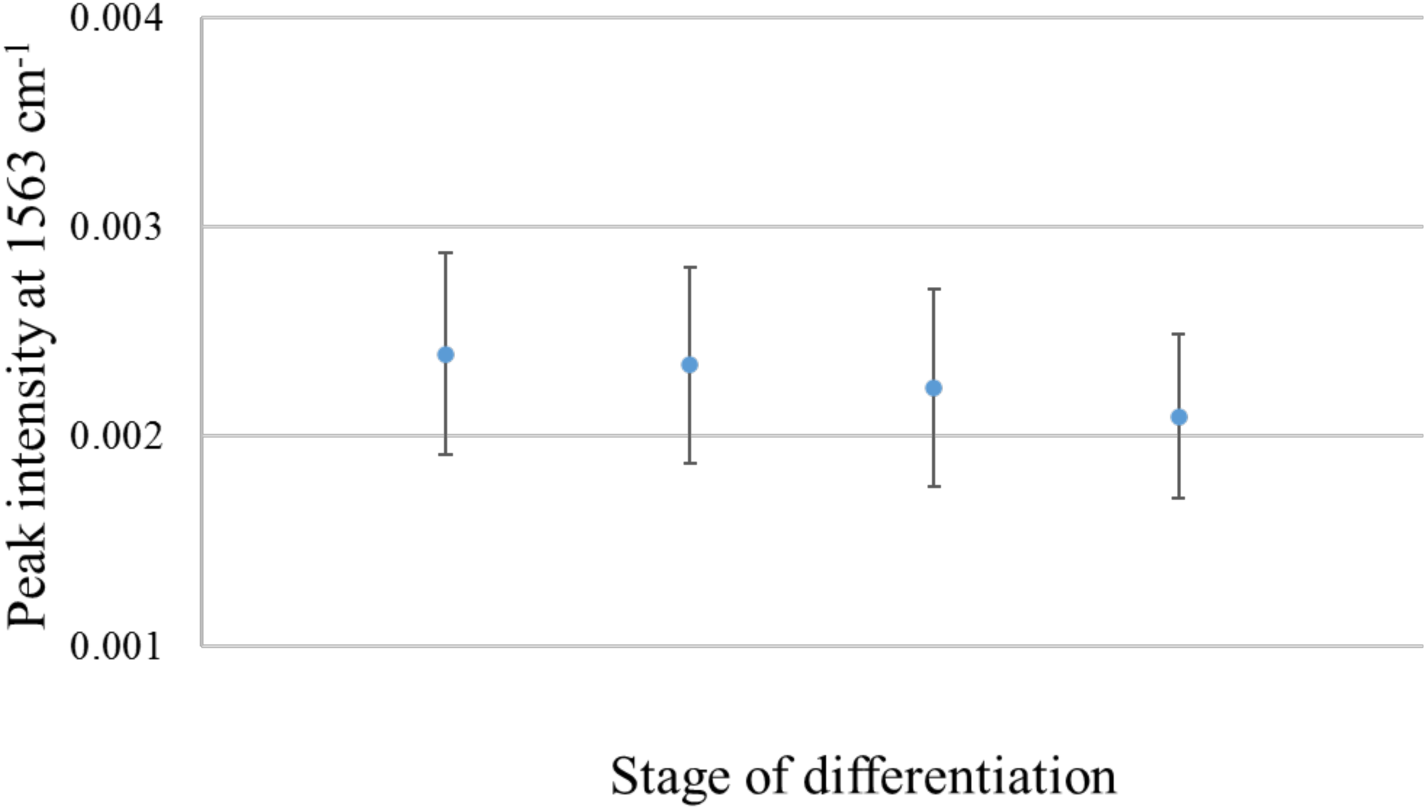
The mean ±1 standard deviation of the 1563 cm^-1^ peak intensity for each of the four classes, demonstrating insignificant differences as observed by the overlap in standard deviations between groups.

**Figure S.3.**
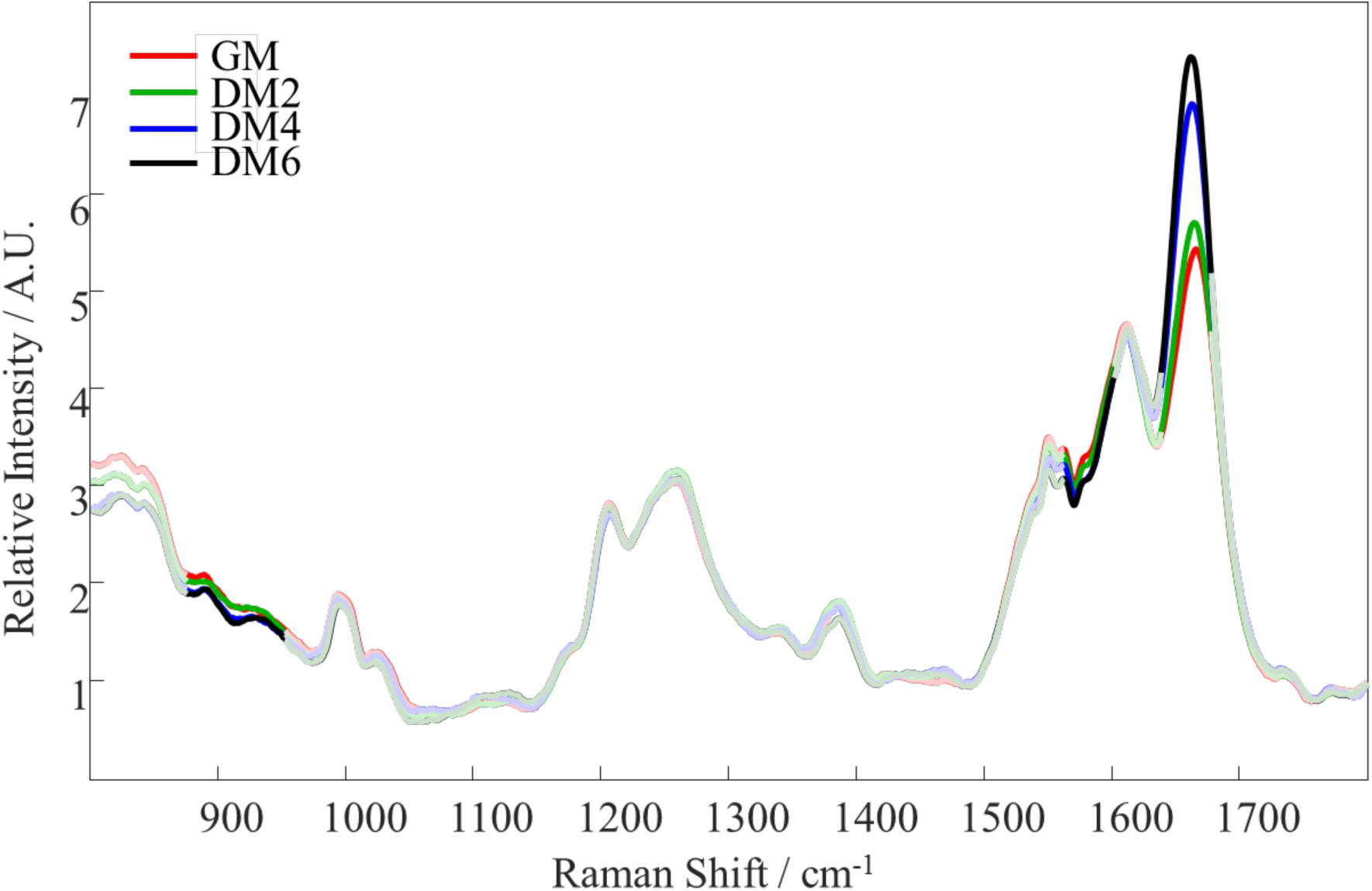
Mean DUVRS spectra of myoblast cell differentiation at various stages, including the spectral ranges selected by genetic algorithm: GM (red), DM2 (green), DM4 (blue), and DM6 (black). Areas selected by genetic algorithm are marked by bolded lines whereas spectral regions deemed uninformative for discrimination are seen as unfilled lines.

**Figure S.4.**
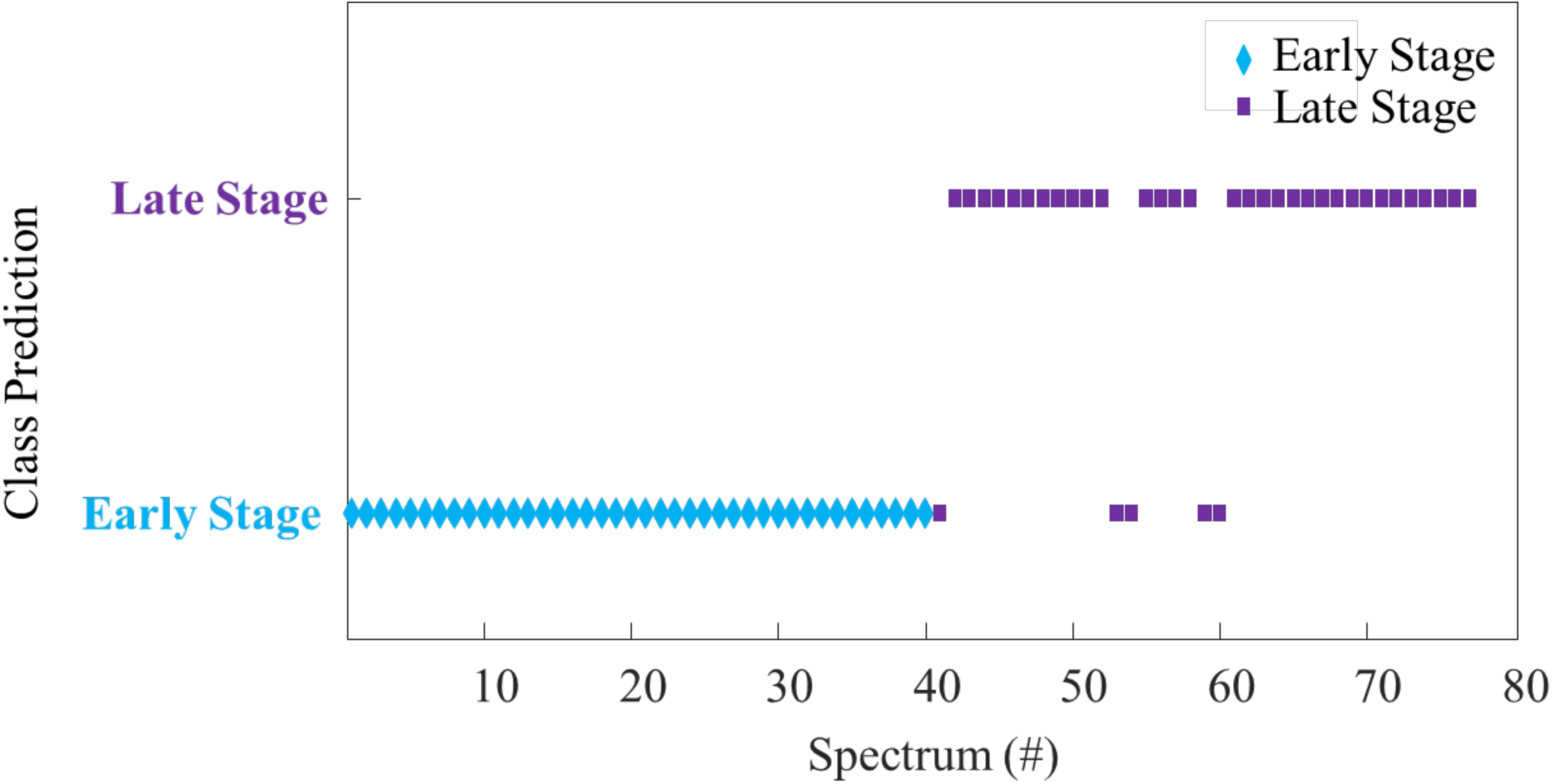
Results of PLS-DA external validation of a binary model for discriminating between early (GM and DM2) and late (DM4 and DM6) stages of myoblast differentiation. Each spectrum from the validation dataset is plotted according to which class it was predicted as belonging: early stage (blue diamond) or late stage (purple square). Each symbol represents one spectrum.

